# Beyond Antibiotics: Cinnamic Acid’s role in combatting complex biofilms in the oral cavity

**DOI:** 10.1101/2024.01.02.573850

**Authors:** Lama Al_Darwish, Muaaz Alajlani, Sharif Alashkar

**Affiliations:** Al_Sham Private University, Faculty of Pharmacy, Damascus, Syria.; Al_Sham Private University, Faculty of Detistry, Damascus, Syria

## Abstract

Traditional antibiotic therapy has become inefficient in treating oral plaques, due to the the presence of highly adherent bacteria in complex infections of oral cavities. Our study aimed to evaluate the anti-biofilm activity of Cinnamic Acid against single and multispecies bacteria causing dental plaques. Different concentrations of cinnamic acid (1-1000 mg/L) were tested using Biofilm assays, EPS analysis, and biomass quantification. To gain insight, a drug-likeness chemoinformatics study of cinnamic acid was conducted.

We found that lower concentrations (1-400mg/L) of Cinnamic Acid had a minor effect on biofilm formation inhibition. Whilst concentrations of 600mg/L and above had shown significant biofilm production, EPS production and biomass production reduced down to 80%.Cinnamic Acid has proven to contain antibacterial, antibiofilm, and drug-likeness properties and is a prominent compound in combating oral biofilm.

**Authors Summary:** Combating oral pathogenic biofilm is an important approach in treating polymicrobial oral infections, We have tested the anti-biofilm activity of cinnamic acid at different connections (1-1000 mg/L) and against single and multispecies biofilm forming bacteria. We found that cinnamic acid had a notable reduction in biofilm formation and EPS production, and reduced total biofilm biomass. Cinnamic Also fulfills the lipinski ROFs criteria for drug-likeness.

Due to its antibacterial, anti-biofilm, and drug-likeness properties, a plant-derived compound such as Cinnamic Acid can be a new prominent approach to defecting the global risk of oral pathogenic biofilms. And serve as an alternative to conventional oral care methods which have been rendered inefficient by biofilm-forming pathogens.

## Introduction

Oral diseases continue to be a major health problem worldwide. Dental plaques, polymicrobial complexes of biofilm-forming microorganisms, represent a challenge in oral health, leading to serious conditions such as Gingivitis, periodontitis, and Systemic Health Implications(1). Furthermore, biofilms play a remarkable role in escaping host immunogenicity(2), making it challenging to eliminate the bacteria within oral infections. As well as its major effect on antibiotic resistance(3), due to the lipophilic adherent nature of EPS. Biofilms, are intricate communities of microorganisms encased in a self-produced extracellular polymeric substance (EPS), To fulfill biological roles such as adhesion, structural scaffolding/cementing, and protection.rendering conventional oral hygiene measures and traditional antibiotic therapy less effective.(4)

The compositions of biofilm matrices differ largely between different bacteria and growth conditions under which biofilms are formed. Characterization of oral biofilms contained Host molecules, Cell wall fragments, Proteins, Extracellular DNA, Teichoic acids, and lipoteichoic acids. As well as an extensive range of carbohydrates such as Protein-linked bacterial glycans: S-layers, outer membrane proteins, flagella, capsular-related polysaccharides, and Poly-*N*-acetyl-D-glucosamine (5).

*Streptococcus mutans* is considered to be the major pathogenic oral biofilm producer causing dental plaques(6). However, studies have shown that Respiratory infection-related bacteria and GIT infection-related bacteria can present in the oral cavity and contribute to oral biofilm formation, including *pseudomonas aeruginosa*, *Escherichia coli*, *Staphylococcus aureus*, and Salmonella spp due to the structural contact between oral, upper gastrointestinal and lower respiratory cavities.(7,8,9)

Nowadays, natural phytochemical antibacterial compounds are attracting attention to develop novel therapeutics against oral infectious diseases(10). Cinnamic Acid(CA). is a phenolic compound found in Cinnamomum zeylanicum, and is a WHO-approved phytochemical known for its antioxidants, anticancer, and antibacterial activity (11).

Our study aims to perform an in vitro and in-sillico evaluation of the anti-biofilm activity of Cinnamic Acid against single and multispecies bacteria causing dental plaques.

## Results

### Biofilm forming ability of bacterial strains

As Shown in (Figure 1). Our study revealed that all bacterial species either single or multispecies, could form a biofilm. However, we found that multispecies, *S. Mutans*, *S.Aures*, and *Pseudomonas spp* had the most prominent potential for biofilm.

**Figure 1:**
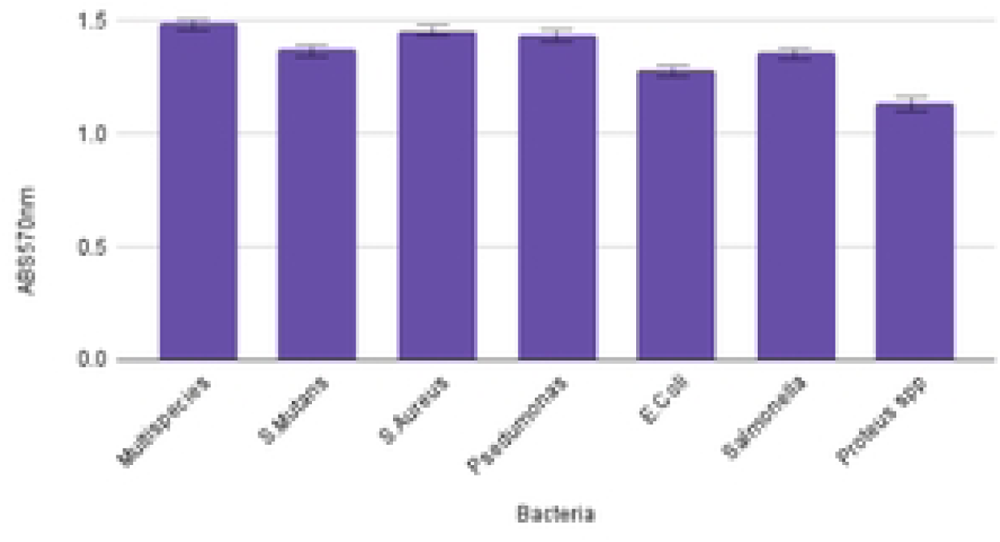
Bipfilm forming ability of bacteria found in dental plauqe.

### Cinnamic Aid effect on biofilm formation in single and multispecies bacteria

The present study indicated that lower concentrations of cinnamic acid(1-400 mg/L) had no or minimal inhibitory effect on biofilm formation in single and multispecies bacteria causing dental plaques. However, a (600-800mg/L)concentration of cinnamic acid has been shown to reduce biofilm formation in single and multispecies bacteria by (46.8-64.7)%. At high concentrations (1000mg/L), Cinnamic Acid has a prominent biofilm inhibition activity of 82.13%. As shown in (Figure 2).

**Figure2:**
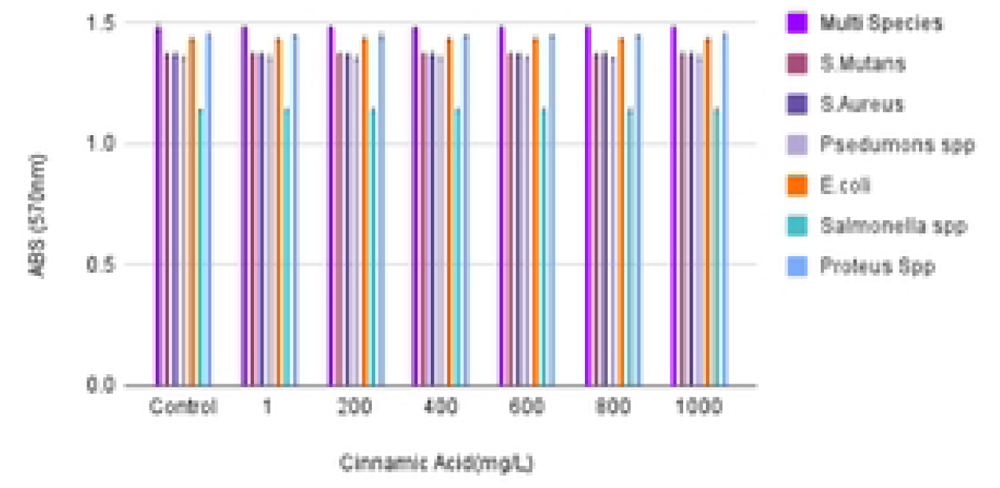
CA **Effect** onbiofilm formation In single and multispecies bacteria.

### Cinnamic Acid effect on extracellular polymeric substance production(EPS) in multispecies bacteria

For the characterization of biofilm production, different concentrations of Cinnamic Acid were analyzed and compared to the untreated sample to check the effects on extra polymeric substance (EPS) production.

Our study presents that high concentrations of C.A(800,1000mg/L)had significantly reduced EPS production to 60% and 70.1%.Compared with other concentrations which had a minor effect on EPS formation inhibition or dispersal. As shown in (Figure 3.)

**Figure 3:**
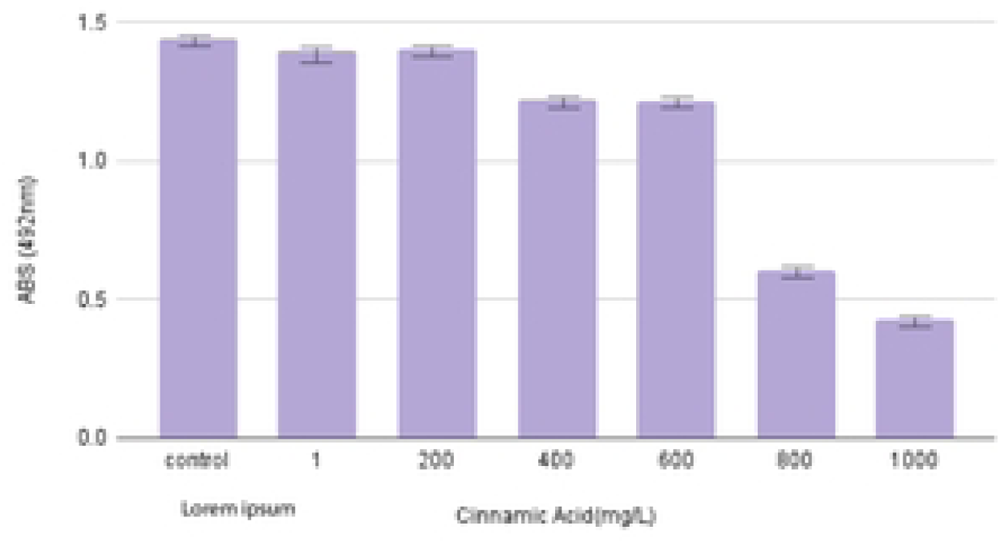
Cinnamic Acid effect on EPS Production.

### Cinnamic Acid Effect on Bacterial Biomass

As shown in(Figure 4), The production of biomass was potentially reduced by applying different concentrations of Cinnamic Acid. Although low concentrations were not able to achieve a high reduction in biomass concentrations nor surface coverage(41%). (1-200mg/L),an extensive biomass reduction at higher (400mg/L and above) Cinnamic Acid concentrations as compared to the control. (up to 88%)

**Figure4:**
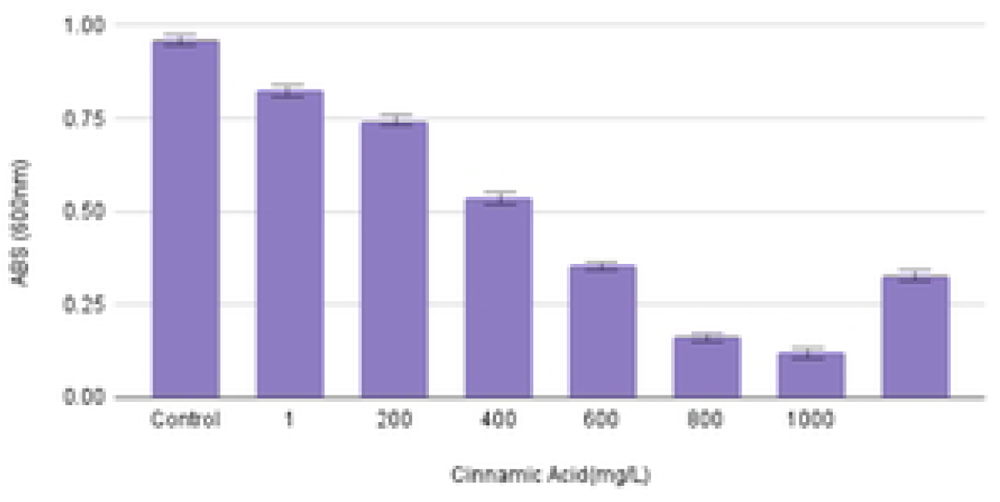
Cinnamic Acid effect on biomass in multispecies bacteria.

### Drug-likeness evaluation of Cinnamic Acid

Depending on the Chemical structure and molecular properties of Cinnamic Acid, the result from chemoinformatic analysis presents an octanol-water partition coefficient (log P) ≤ 5, a molecular weight ≤ 500 Da (g/mol), a number of hydrogen bond acceptors ≤ 10, and a number of hydrogen bond donors ≤ 5, number of rotatable bonds (n-ROTB) ≤ 5 and a topological polar surface area <40 Å. Figure 5.

**Figure5:**
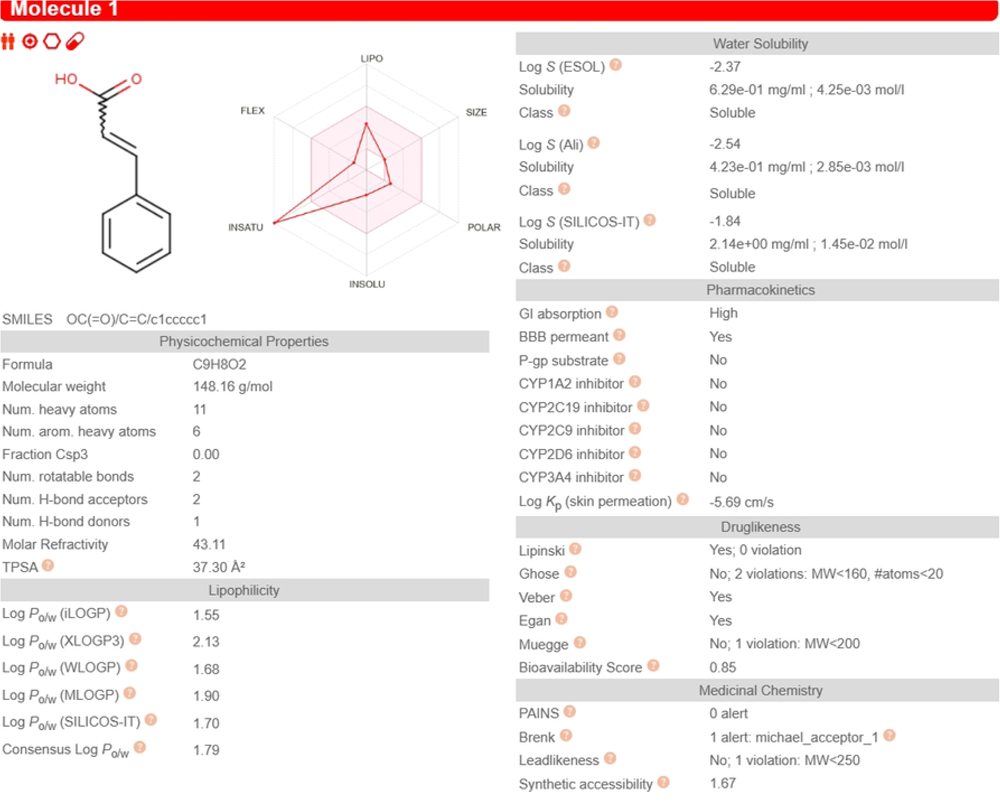

This indicates that Cinnamic Acid fulfills the Lipinski rule of five criteria to be considered a drug-like compound.(12)

## Discussion

Uncontrolled oral biofilm resulting from pathogenic and carcinogenic bacteria Proteus spp., Escherichia coli, Pseudomonas spp., Salmonella spp., Streptococcus mutans., and Staphylococcus aureus can cause serious systemic health hazards.(13)

Our study aimed to investigate the prevalence of antibiofilm properties in cinnamic acid and to investigate the dose-response relationship of CA using a range of low and high concentrations(1-1000 mg/L). We tested the biofilm-forming potential of a single and multispecies bacteria isolated from a polymicrobial infectious dental plaque. Next, We performed a biofilm assay, EPS assay, and Biomass assay after applying different concentrations of CA. Multiple researchers have used this similar protocol to test the antibiofilm activity of phenolic compounds(14), such as (Albutti, Aet al) who tested Gallic Acid Antibiofilm and biofilm dispersal properties against *Proteus spp*., *Escherichia coli*, *Pseudomonas spp*., *Salmonella spp*., *Streptococcus mutans*., and *Staphylococcus aureus* as well as multispecies bacteria(15). Also,(<u>Lin Yue</u> <u>et al</u>) have tested Cinnamic Acid derivatives as antibiofilm agents against multispecies bacteria(16). Another interesting study by <u>(Hongmei Zhang</u> Et al) have documented the inhibitory effect of CA against biofilm formation in *Enterobacteriaceae* (17).

We found that a concentration of 1mg/L, 200mg/L, and 400mg/L cinnamic reduced biofilms concentration by 6.2%, 20.57%, and 33.18%.Applying 600mg/L, 800mg/L, and 1000mg/L CA resulted in significant biomass production inhibition by 46.8%, 65%, and 82.2% in single and multispecies bacteria.

We suggest that CA was able to inhibit biofilm production by interfering with the biological mechanism of biofilm forming. In *Enterobacteriaceae*, Curli amyloid fibrils mediate host cell adhesion and contribute to biofilm formation, thereby promoting bacterial resistance to environmental stressors(18). Structural insights into Crurli fibrils have revealed that major and minor subunits CsgA and CsgB, and chaperon-like protein CsgC, are encoded transcription of the csgBAC operon, and under controlled by CsgD(19). Our study suggests an interaction between CA and major biofilm regulator CsD. Which may be the cause of the antibiofilm activity of CA against *Escherichia coli* and *Sallonmnella spp.* strains. In streptococcus mutans, extracellular Glucosyltransferases are critical for the synthesis of homopolymers Glucan by the use of glucose(20). We present that CA can bind to Glucosetransferase and inhibit its action. Which we suggest is the cause of CA antibiofilm activity against *Streptococcus mutans*.(21).

Type IV pili are expressed by opportunistic bacteria like *Pseudomonas spp*, Including PilY1 and PilY2 have been associated with biofilm formation and cell adhesion. We suggest that cinnamic acid formed a bond with the PilY1 C-terminal domain and we correlate this result with the in vitro antibiofilm assessment of CA.(22,23)

We also suggest a connection between the antibiofilm property of CA against S.Aures by binding to the C domain of *staphylococcal* protein A mutant and EAP Domains(24).

Since Eps is the major biofilm matrice component related to cell adhesion (25), we tested different concentrations of CA against EPS Production in multispecies. We found that a high concentration of CA (800mg/L and above) is a prominent agent in the reduction of EPS formation in multispecies (57.5% to 70%). Higher concentrations of CA have also resulted in a notable reduction in Biomass down to 83%.

We think these results have emerged due to the biofilm production inhibition property of CA. Suggesting that CA has a biofilm dispersal activity alongside inhibiting its formation.

We have conducted a chemoinformatics study to suggest a drug-likeness and ADMET model of CA depending on the Lipinski rule of five. The lipophilic nature of Cinnamic Acid(Logp<5) allows it to interact with the thick cell membrane of gram-positive bacteria. Whilst a TPSA<40 can cause the interaction with Gram Negative bacteria(26) Furthermore, Mw below 500g/mol, Hbd<5 and HBA<10, Logp<5, and TPSA<40 are complied with the ROFs suggest that CA is a drug like compound. (27)

## Materials and Methodology

### Sample Collection

The participant in this study was 49 male, who presented to the dental clinic with multiple plaques on the gingivae, inside of the cheeks, and tongue persisting for over 14 days.

Our participant had poor dental status, a medical history of extensive dental treatment within the past year, and reported an irrational use of antibiotics. The exclusion criteria were patients with a stable oral microbiome.

The reasoning for this criteria is to capture a complex microbial community, reflecting the real-world scenario of polymicrobial biofilms.

The biofilm isolation procedure was performed with the assistance of a licensed dentist and included gentle scraping of the lesions using a sterile cell scraper.

Written informed consent was obtained from the participant, and ethical approval for the study has been granted from the scientific deanship of Al-Sham Private University.

### Bacterial Strains Isolation and Identification

After isolation from the oral cavity, the biofilms were transferred to Eppendorf tubes containing 2 mL sterile Tryptic Soy Broth supplemented with 2 mL sterile sucrose solution 1%, incubated for 24 hours at 37 C to enhance bacterial growth.

The suspension was inoculated on MacConkey Agar, Eosin Methylene Blue (EMB) Agar, Cetrimide agar, Xylose Lysine Deoxycholate (XLD) Agar, Mannitol Salt Agar (MSA) and Blood Agar, and Incubated for 48hours at 36 ± 1.

Depending on the common oropharyngeal microbiota as described by the European Society of Clinical Microbiology and Infectious Diseases,(28). Bacterial strains that are not part of the common oropharyngeal microbiota were considered potential pathogens.

Six different dental plaque bacterial species were isolated and identified, including *Proteus spp*., *Escherichia coli*, *Pseudomonas spp*., *Salmonella spp*., *Streptococcus mutans.*, and *Staphylococcus aureus*. (29)

Experimental conditions, including bacterial strains, growth media, and incubation parameters, were standardized across all in vitro experiments.

### Biofilm Crystal Violet Assay

To test their biofilm-forming properties, single and multispecies bacteria were grown in 24-well microtiter plates, each tube containing 250µl of bacterial suspension prepared from the initial culture (adjusted to 0.5 on Mc Farland turbidity) maintained in 1 mL of sterile Tryptic Soy Broth supplemented with sterile sucrose solution 1%, incubated for 24 hours at 37C.

After incubation, the wells were emptied, and washed with 1X PBS sterile Phosphate Buffer Saline Solution (PH=7.2) to remove excess media and planktonic cells.

The remaining adherent cells were fixed using 250µl methanol96% per well for 10 mins, stained with 200µl crystal violet dye 0.1% per well for 10 min at room temperature.

The excess stain was washed using sterile distilled water. Then, excess liquid was poured off and the plates were left to air dry for 5 minutes.

The dye bound to the adherent cells was re-dissolved with 200 µl of 33% (v/v) glacial acetic acid, moved to cuvettes, and quantified by measuring the optical Absorbance at 570 nm using a spectrophotometer. (30.31)

For positive and negative control, 9 wells contained Sterile Tryptic Soy Broth.

### Biofilm Calculations

Bacterial strains were classified into different categories depending on their obtained optical density.(32)

Nonadherent: OD < ODC.

Weakly adherent: ODC < OD < 2ODC

Moderately adherent : 2ODC < OD < 4ODC

strong biofilm producer: 4ODC < OD

### Cinnamic Acid Treatment

Trans (E) Extra pure 99% Cinnamic Acid, purchased from titan-biotech in the form of crystal powder, was used in this experiment. Different concentrations (1,200,400,600,800, 1000 mg/L) of Cinnamic Acid were tested for their biofilm inhibition ability, and were prepared accordingly:

Powdered cinnamic Acid was weighed accurately, moved to a sterile glass container, carefully dissolved in dimethyl sulfoxide DMSO1%, and sterile distilled water at room temperature.

The wells were labeled for each concentration of Cinnamic Acid in triplicate (1,200,400,600, 800, 1000 mg/L). Instead of Cinnamic Acid, 50 µL of sterilized distilled water was added to the positive and negative control wells.

### Anti-Adherent Activity Assay (Anti Biofilm Crystal Violet Assay)

This assay was performed according to the microtiter method previously described(30). Pre-formed biofilms (48 hours old) were used for this assay. The established biofilms were washed three times with sterile PBS solution, and then 50µl of Cinnamic Acid (1–1000 mg/L) solution was exposed for 15mins.

After that, the biofilm was quantified by the crystal violet assay. For Blank(Untreated) Broth was considered.

### Extracellular Polymeric Substance Assay

Only multispecies bacteria were examined in this assay, Multispecies were grown on glass slides surface in a petri dish in a nutrient broth medium, by adding 500µl of bacterial suspension to 1 mL of nutrient broth and maintaining the samples at 37 C for 48 hours.

After incubation, 50µl of CA (1–1000mg/L) solution was applied and left for 24 hours at 37C.

Control was considered to be 50µl of sterile distilled water.

After that, EPS was extracted using a cell scrapper and transferred to a sterile glass tube containing 5 mL of PBS solution. The sample was mixed in a vortex for 40 sec. Then, all tubes were centrifuged using a centrifuge machine at room temperature for 15 min at 10,000 rpm.

The supernatant material was transferred into a fresh test tube and was considered to include soluble EPS. The pellets down the tube’s bottom were considered to be cell biomass.

1 mL of the soluble EPS was poured into a sterile glass tube containing 500µl of phenol 5%. Then, 2 mL of concentrated sulfuric acid solution was added carefully to the walls of the tube. The mixture was incubated for 10 minutes at room temperature.

Finally, the mixture was transferred to a quartz cuvette and the EPS was quantified at 492 nm using a UV spectrophotometer.(33)

### Biomass concentration determination

Only multispecies bacteria were examined in this assay, previously centrifuged samples were used, and pellets down the tube’s bottom were considered to be cell biomass.

After EPS Extraction, the centrifuged tubes containing pellets were washed with 10 mL saline, and 5mL PBS was added. The samples were further mixed by a vortex for 1 minute a. Then, their optical Absorbance was obtained using a spectrophotometer at 600nm.(34)

### Drug Likness Evaluation

Using SWISS ADMET Software <u>SwissADME</u>, molecular and physiochemical properties of CA were prepared.We used the lipiniski rule of fives as a measurement for drug likness.

### Reagents and Chemicals

Cinnamic Acid was purchased from Titan Biotech, India.

DMSO, Ringer Solution, PBS Solution, and Sulfuric Acid, Crystal violet stain, powdered was purchased from Sigma Aldrich.

All tubes, dishes, cuvettes, and microtiter plates were purchased from a local market.

### Statistical Analysis

All experiments were performed in triplicates, Statistical analyses were performed using IBM SPSS Statistics version 26 (Chicago, IL, USA) and Microsoft Excel 10 (Microsoft, Redmond, WA, USA). For all of the hypotheses tested, a p-value of less than 0.05 indicated statistical significance

## Acknowledgments

None.

## Supporting information

**S1 Fig.:** Biofilm Forming ability of bacterial strains isolated from oral dental plauqes.

**S2 Fig:** Cinnamic Acid Effect on biofilm formation in singl and multispecies bacteria in microtiter pates.

**S3 Fig:** Cinnamic Acid effect on EPS production in multispecies.

**S4 Fig:** Cinnamic Acid effect on biomass in multispecies.

## References

1. Christersson, L. A., Zambon, J. J., & Genco, R. J. (1991). Dental bacterial plaques. Nature and role in periodontal disease. Journal of clinical periodontology, 18(6), 441–446. 10.1111/j.1600-051x.1991.tb02314.x

2. Yamada, K. J., & Kielian, T. (2019). Biofilm-Leukocyte Cross-Talk: Impact on Immune Polarization and Immunometabolism. Journal of innate immunity, 11(3), 280–288. 10.1159/000492680

3. Rather, M. A., Gupta, K., & Mandal, M. (2021). Microbial biofilm: formation, architecture, antibiotic resistance, and control strategies. Brazilian journal of microbiology : [publication of the Brazilian Society for Microbiology], 52(4), 1701– 1718. 10.1007/s42770-021-00624-x

4. O’Toole, G., Kaplan, H. B., & Kolter, R. (2000). Biofilm formation as microbial development. Annual review of microbiology, 54, 49–79. 10.1146/annurev.micro.54.1.49

5. Jakubovics, N. S., Goodman, S. D., Mashburn-Warren, L., Stafford, G. P., & Cieplik, F. (2021). The dental plaque biofilm matrix. Periodontology 2000, 86(1), 32–56. 10.1111/prd.12361

6. Lemos, J. A., Palmer, S. R., Zeng, L., Wen, Z. T., Kajfasz, J. K., Freires, I. A., Abranches, J., & Brady, L. J. (2019). The Biology of *Streptococcus mutans*. Microbiology spectrum, 7(1), 10.1128/microbiolspec.GPP3-0051-2018.

7. El-Telbany, M., & El-Sharaki, A. (2022). Antibacterial and anti-biofilm activity of silver nanoparticles on multi-drug resistance *pseudomonas aeruginosa* isolated from dental-implant. Journal of oral biology and craniofacial research, 12(1), 199–203. 10.1016/j.jobcr.2021.12.002

8. Perkowski, K., Baltaza, W., Conn, D. B., Marczyńska-Stolarek, M., & Chomicz, L. (2019). Examination of oral biofilm microbiota in patients using fixed orthodontic appliances in order to prevent risk factors for health complications. Annals of agricultural and environmental medicine : AAEM, 26(2), 231–235. 10.26444/aaem/105797

9. Schnurr, E., Paqué, P. N., Attin, T., Nanni, P., Grossmann, J., Holtfreter, S., Bröker, B. M., Kohler, C., Diep, B. A., Ribeiro, A. A., & Thurnheer, T. (2021). *Staphylococcus aureus* Interferes with Streptococci Spatial Distribution and with Protein Expression of Species within a Polymicrobial Oral Biofilm. *Antibiotics (Basel*, Switzerland), 10(2), 116. 10.3390/antibiotics10020116

10. Ribeiro, M., Malheiro, J., Grenho, L., Fernandes, M. H., & Simões, M. (2018). Cytotoxicity and antimicrobial action of selected phytochemicals against planktonic and sessile *Streptococcus mutans*. PeerJ, 6, e4872. 10.7717/peerj.4872

11. Upadhyay, A., Upadhyaya, I., Kollanoor-Johny, A., & Venkitanarayanan, K. (2014). Combating pathogenic microorganisms using plant-derived antimicrobials: a minireview of the mechanistic basis. BioMed research international, 2014, 761741. 10.1155/2014/761741

12. Lipinski, C. A., Lombardo, F., Dominy, B. W., & Feeney, P. J. (2001). Experimental and computational approaches to estimate solubility and permeability in drug discovery and development settings. Advanced drug delivery reviews, 46(1-3), 3–26. 10.1016/s0169-409x(00)00129-0

13. Valm A. M. (2019). The Structure of Dental Plaque Microbial Communities in the Transition from Health to Dental Caries and Periodontal Disease. Journal of molecular biology, 431(16), 2957–2969. 10.1016/j.jmb.2019.05.016

14. Darmasiwi, S., Aramsirirujiwet, Y., & Kimkong, I. (2022). Antibiofilm activity and bioactive phenolic compounds of ethanol extract from the *Hericium erinaceus* basidiome. Journal of advanced pharmaceutical technology & research, 13(2), 111–116. 10.4103/japtr.japtr_1_22

15. Albutti, A., Gul, M. S., Siddiqui, M. F., Maqbool, F., Adnan, F., Ullah, I., Rahman, Z., Qayyum, S., Shah, M. A., & Salman, M. (2021). Combating Biofilm by Targeting Its Formation and Dispersal Using Gallic Acid against Single and Multispecies Bacteria Causing Dental Plaque. *Pathogens (Basel*, Switzerland), 10(11), 1486. 10.3390/pathogens10111486

16. Yue, L., Wang, M., Khan, I. M., Xu, J., Peng, C., & Wang, Z. (2021). Preparation, characterization, and antibiofilm activity of cinnamic acid conjugated hydroxypropyl chitosan derivatives. International journal of biological macromolecules, 189, 657–667. 10.1016/j.ijbiomac.2021.08.164

17. Zhang, H., Zhou, W., Zhang, W., Yang, A., Liu, Y., Jiang, Y., Huang, S., & Su, J. (2014). inhibitory effects of citral, cinnamaldehyde, and tea polyphenols on mixed biofilm formation by foodborne Staphylococcus aureus and Salmonella enteritidis. Journal of food protection, 77(6), 927–933. 10.4315/0362-028X.JFP-13-497

18. Serra, D. O., & Hengge, R. (2021). Bacterial Multicellularity: The Biology of *Escherichia coli* Building Large-Scale Biofilm Communities. Annual review of microbiology, 75, 269–290. 10.1146/annurev-micro-031921-055801

19. Perov, S., Lidor, O., Salinas, N., Golan, N., Tayeb-Fligelman, E., Deshmukh, M., Willbold, D., & Landau, M. (2019). Structural Insights into Curli CsgA Cross-β Fibril Architecture Inspire Repurposing of Anti-amyloid Compounds as Anti-biofilm Agents. PLoS pathogens, 15(8), e1007978. 10.1371/journal.ppat.1007978

20. Ren, Z., Cui, T., Zeng, J., Chen, L., Zhang, W., Xu, X., Cheng, L., Li, M., Li, J., Zhou, X., & Li, Y. (2015). Molecule Targeting Glucosyltransferase Inhibits Streptococcus mutans Biofilm Formation and Virulence. Antimicrobial agents and chemotherapy, 60(1), 126–135. 10.1128/AAC.00919-15

21. Laverty, G., Gorman, S. P., & Gilmore, B. F. (2014). Biomolecular Mechanisms of Pseudomonas aeruginosa and Escherichia coli Biofilm Formation. *Pathogens (Basel*, Switzerland), 3(3), 596–632. 10.3390/pathogens3030596

22. Marko, V. A., Kilmury, S. L. N., MacNeil, L. T., & Burrows, L. L. (2018). Pseudomonas aeruginosa type IV minor pilins and PilY1 regulate virulence by modulating FimS-AlgR activity. PLoS pathogens, 14(5), e1007074. 10.1371/journal.ppat.1007074

23. Speziale, P., Pietrocola, G., Foster, T. J., & Geoghegan, J. A. (2014). Protein-based biofilm matrices in Staphylococci. Frontiers in cellular and infection microbiology, 4, 171. 10.3389/fcimb.2014.00171

24. Zhang, R., Neu, T. R., Blanchard, V., Vera, M., & Sand, W. (2019). Biofilm dynamics and EPS production of a thermoacidophilic bioleaching archaeon. New biotechnology, 51, 21–30. 10.1016/j.nbt.2019.02.002

25. Zhang, R., Neu, T. R., Blanchard, V., Vera, M., & Sand, W. (2019). Biofilm dynamics and EPS production of a thermoacidophilic bioleaching archaeon. New biotechnology, 51, 21–30. 10.1016/j.nbt.2019.02.002

26. Tommasi, R., Brown, D. G., Walkup, G. K., Manchester, J. I., & Miller, A. A. (2015). ESKAPEing the labyrinth of antibacterial discovery. Nature reviews. Drug discovery, 14(8), 529–542. 10.1038/nrd4572

27. Walters W. P. (2012). Going further than Lipinski’s rule in drug design. Expert opinion on drug discovery, 7(2), 99–107. 10.1517/17460441.2012.648612

28. Ewan VC, Reid WDK, Shirley M, Simpson AJ, Rushton SP, Wade WG. Oropharyngeal Microbiota in Frail Older Patients Unaffected by Time in Hospital. Front Cell Infect Microbiol. 2018 Feb 20;8:42. doi: 10.3389/fcimb.2018.00042. PMID: 29515974; PMCID: PMC5826060.

29. Khalid, M., Hassani, D., Bilal, M., Butt, Z. A., Hamayun, M., Ahmad, A., Huang, D., & Hussain, A. (2017). Identification of oral cavity biofilm forming bacteria and determination of their growth inhibition by Acacia arabica, Tamarix aphylla L. and Melia azedarach L. medicinal plants. Archives of oral biology, 81, 175–185. 10.1016/j.archoralbio.2017.05.011

30. Tahmourespour, A., Aminzadeh, A., & Salehifard, I. (2022). Anti-adherence and anti-bacterial activities of *Pistacia atlantica* resin extract against strongly adherent *Streptococcus mutans* strains. Dental research journal, 19, 36.

31. Kragh, K. N., Alhede, M., Kvich, L., & Bjarnsholt, T. (2019). Into the well-A close look at the complex structures of a microtiter biofilm and the crystal violet assay. Biofilm, 1, 100006. 10.1016/j.bioflm.2019.100006

32. Stepanović, S., Vuković, D., Hola, V., Di Bonaventura, G., Djukić, S., Cirković, I., & Ruzicka, F. (2007). Quantification of biofilm in microtiter plates: overview of testing conditions and practical recommendations for assessment of biofilm production by staphylococci. APMIS : acta pathologica, microbiologica, et immunologica Scandinavica, 115(8), 891–899. 10.1111/j.1600-0463.2007.apm_630.x

33. Di Martino P. (2018). Extracellular polymeric substances, a key element in understanding biofilm phenotype. AIMS microbiology, 4(2), 274–288. 10.3934/microbiol.2018.2.274

34. Peeters, E., Nelis, H. J., & Coenye, T. (2008). Comparison of multiple methods for quantification of microbial biofilms grown in microtiter plates. Journal of microbiological methods, 72(2), 157–165. 10.1016/j.mimet.2007.11.010

